# Ultra high diversity factorizable libraries for efficient therapeutic discovery

**DOI:** 10.1101/2022.01.17.476670

**Authors:** Zheng Dai, Sachit D. Saksena, Geraldine Horny, Christine Banholzer, Stefan Ewert, David K. Gifford

**Author notes:** Equal contribution (Zheng Dai, Sachit D. Saksena).

## Abstract

The successful discovery of novel biological therapeutics by selection requires highly diverse libraries of candidate sequences that contain a high proportion of desirable candidates. Here we propose the use of computationally designed factorizable libraries made of concatenated segment libraries as a method of creating large libraries that meet an objective function at low cost. We show that factorizable libraries can be designed efficiently by representing objective functions that describe sequence optimality as an inner product of feature vectors, which we use to design an optimization method we call Stochastically Annealed Product Spaces (SAPS). We then use this approach to design diverse and efficient libraries of antibody CDR-H3 sequences with various optimized characteristics.

## 1 Introduction

Biologics, such as monoclonal antibody therapeutics, are commonly discovered by expressing diverse candidate libraries in phage or yeast display systems followed by multiple rounds of affinity selection against a biological target of interest [12]. Many other protein engineering tasks, including discovery of adeno-associated vectors (AAV) for gene therapy [19, 1], T-cell receptor (TCR) design [4, 10, 18], and aptamer screening [13, 8] can also be framed as selection from a library of candidates.

Therapeutic discovery by selection requires candidate libraries that are both highly diverse and enriched in desirable candidates in order to isolate even a single lead for further pre-clinical development. We define *library diversity* as the number of sequences in a library that are sufficiently different from each other to produce different therapeutics. We define *library efficiency* as the proportion of library sequences with favorable therapeutic, delivery, and manufacturing properties [14]. State-of-the-art antibody candidate libraries are typically produced via randomization with experimental methods such as trinucleotide (codon) synthesis or mutagenesis, which produce highly diverse libraries that prioritize exploration of the possible sequence space [16]. However, these methods produce inefficient libraries that contain sequences with undesirable qualities such as polyspecificity, hydrophobicity, and instability. Such properties have negative consequences that range from manufacturing difficulty to dangerous clinical side-effects [15]. In recent years, *designed* libraries where each sequence is explicitly specified have been proposed. Sequences in designed libraries can have superior developability profiles, resulting in efficient libraries [11, 17]. However these designed libraries can be both computationally intensive to design and costly to manufacture. At present the cost of designed libraries is prohibitive for library complexities above ~ 10^6^ sequences, and thus designed libraries are typically not sufficiently diverse for *de novo* therapeutic discovery [6].

Here, we introduce *factorizable libraries* where each library member is a combination of *segments* where each *segment library* is much less complex than a resulting factorizable library. To create a factorizable library, segment libraries are combined, inspired in part by the natural use of recombination to create highly diverse natural libraries of antibodies and T-cell receptors. Importantly, this factorization allows for the synthesis of segment libraries at a low-cost that when combined result in a high-complexity library with desirable properties. We develop a method for designing factorizable libraries efficiently, which we call Stochastically Annealed Product Spaces (SAPS). SAPS iteratively improves segment libraries with respect to an objective function that evaluates the result of their concatenation into a full length factorizable library. After the synthesis of segment library DNA oligonucleotides, segment libraries can be joined with a combination of overhang and blunt end ligation similar to Golden Gate Assembly to create a factorizable library [2, 3].

We demonstrate the utility of this appraoch by designing factorizable libraries that randomize the third complementarity determining region of antibody heavy chains (CDR-H3s). We first show that scoring functions compatible with SAPS can successfully capture experimental measurements of CDR-H3 affinity as measured by phage display and affinity selection experiments conducted with experimentally randomized libraries. Next, we show that this framework can generate factorized CDR-H3 libraries that, when joined combinatorially, contain 10^9^ unique sequences with highly specific and flexible design parameters.

## 2 Methods

We introduce a method for designing segment libraries that when joined create a factorizable library that is optimized for a specific performance objective. The performance objective consists of an efficiency term and a diversity (entropy) term. Here we focus on factorizable libraries that consist of two segments, a prefix segment and a suffix segment. We first show that the task of optimizing a factorizable library, when formalized, is an NP complete problem. We then make the segment library design problem tractable by using simulated annealing as a heuristic meta-algorithm (Figure 1A). Simulated annealing requires multiple evaluations of the objective function to generate distributions of proposed updates, which is prohibitively expensive if computing the objective function requires scoring every sequence. We speed up this evaluation by expressing our objective function as the inner product of features of the prefix segment and features of the suffix segment. This objective formulation allows us to rapidly evaluate the objective by keeping a running sum of these features. For added sequence diversity we also include an entropy term in the optimization objective (Figure 1B). The result of library design is the set of sequences in the prefix library and the set of sequences in the suffix library. Joining all prefix segments with all suffix segments combinatorially yields a factorizable library that realizes the provided objective.

**Fig. 1.**
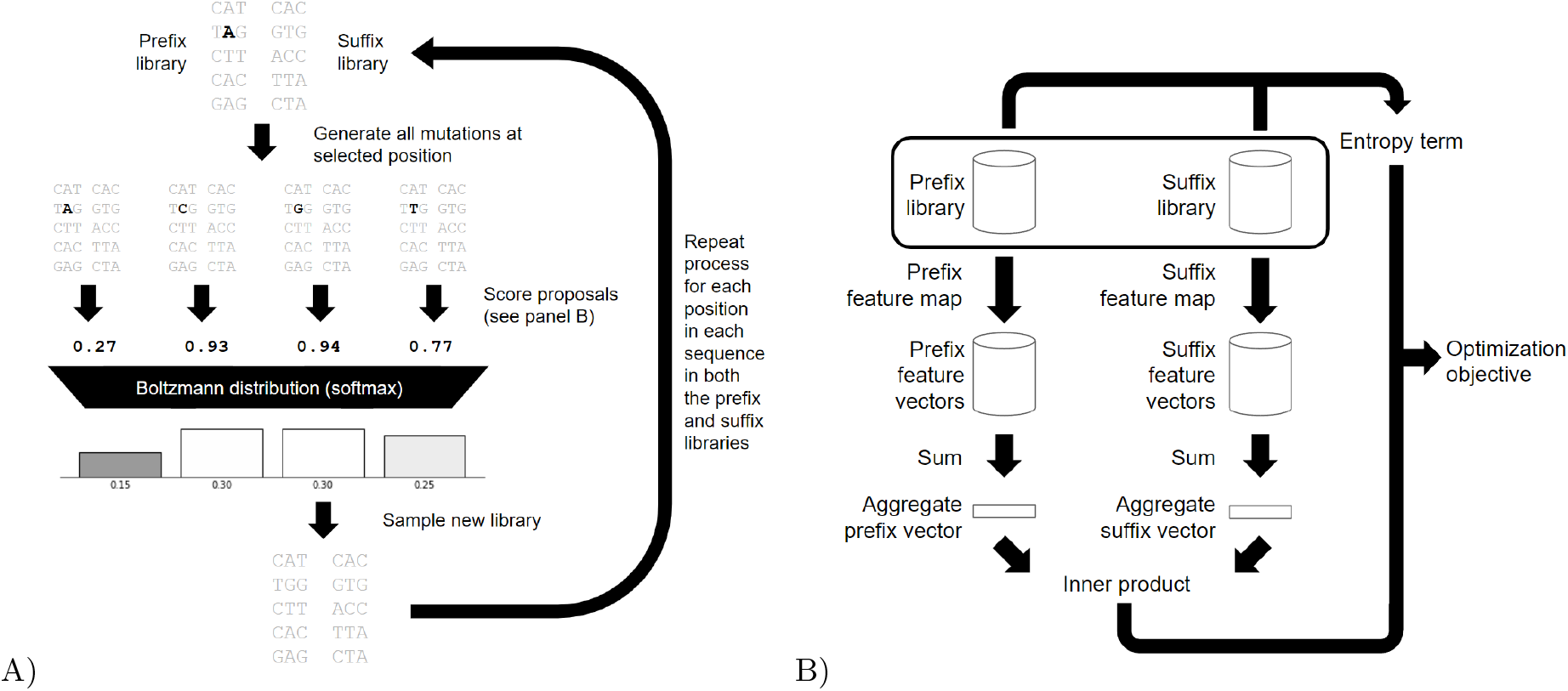
Factorizable library evaluation and optimization. A) Optimization is achieved through iterative stochastic updates. An update step is performed by selecting a position in a sequence in one of the libraries and generating all possible mutations for that position. The mutated libraries are then scored, and then a Boltzmann distribution over the libraries is generated using the negated scores as energy values. The update is then sampled from the distribution. A full update sweep performs this for all positions in all sequences in both libraries. Multiple sweeps are done and the temperature of the Boltzmann distribution is lowered over time. For simplicity, the figure depicts this optimization on small DNA libraries. B) The score of the factorizable library is evaluated by mapping all the sequences in its prefix and suffix libraries to feature spaces. The feature vectors are then aggregated, and an inner product is taken between them, which by the distributive property produces the total score for the whole factorizable library. A position based entropy term is evaluated to quantify the diversity of sequences in the library, and a weighted sum of the two is then used to guide optimization.

### 2.1 Preliminaries

Fix *Σ* to be a finite alphabet and fix *L* to be a positive integer. Let *Σ*^L^ denote the set of strings of length *L* whose symbols are in *Σ*. We also fix two positive integers *L*_p_ and *L*_s_ such that *L*_p_ + *L*_s_ = *L*. We will call *Σ*^L^ the sequence space, 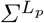 the prefix space, and 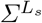 the suffix space.

For a pair of strings *x* and *y*, we use *x* · *y* to denote their concatenation. If *X* and *Y* are instead sets of strings, then their concatenation *X* · *Y* is defined to be {*x* · *y*|*x* ∈ *X, y* ∈ *Y*}.

We use 〈*x, y*〉 to denote the inner product of *x* and *y*, and we will use *x· y* to explicitly denote a dot product if *x* and *y* belong to some Euclidean space. We will use 2^*X*^ to denote the power set of *X*.

We will state various theorems as we explain our method. The proofs are omitted from the main text for improved flow and can be found in Appendix A.

### 2.2 Designing a factorizable library is computationally difficult

Our goal is to design a *factorizable* library of sequences that is *efficient*. To formalize this, suppose we are given some scoring function *f* : *Σ*^L^ → ℝ that characterizes the utility of a single sequence, and we aLre given a pair of positive integers *n*_*p*_ and *n*_*s*_. Then the goal is to find a set *S* ⊆ *Σ*^L^ such that the total score Σ_*s*∈*S*_ *f*(*s*) is maximized, subject to the constraint that *S* is the concatenation of 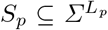 and 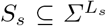 such that |*S*_p_| = *n*_p_ and |*S*_s_| = *n*_s_.

The factorizable constraint adds an additional layer of complexity to the problem, making it much more difficult than library design without the constraint. Supposing that *f*(.) can be expressed as some multi-output boolean circuit of size at most |*Σ*^L^|, then we can always maximize the total score of a library by brute force enumeration in time polynomial in |*Σ*^L^|. However, once factorizability is enforced, then under the assumption that P ≠ NP there exists no algorithm that runs in time polynomial in |*Σ*^L^| that can yield a solution appreciably better than a random solution (Theorem 1 below). Thus, even brute force enumeration of the sequence landscape can be insufficient. This is due to the problem’s relation to finding a large clique in a graph, which is believed to be computationally hard [20].

**Fig. 2.**
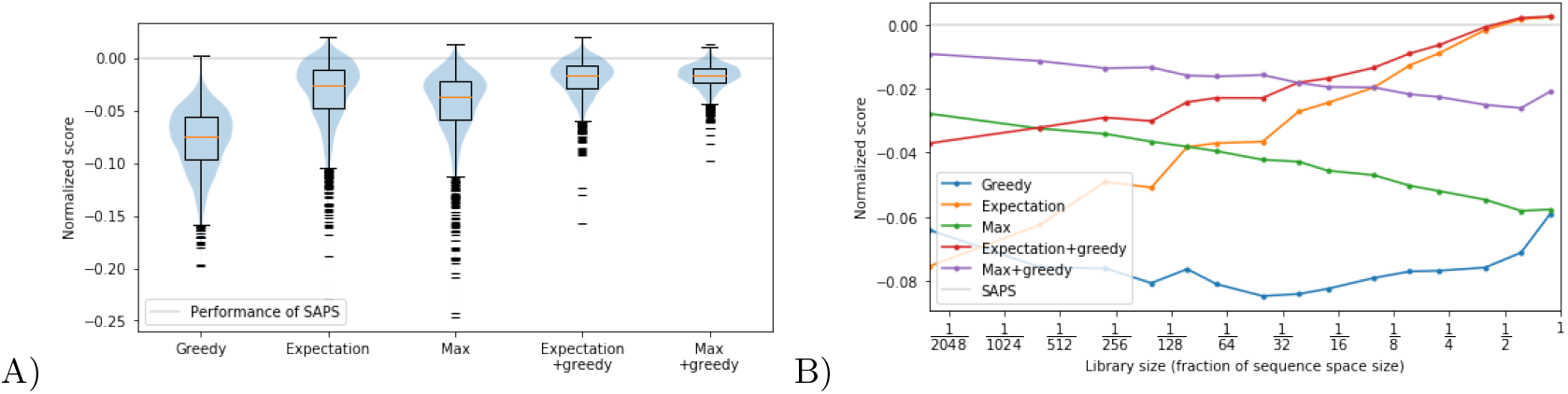
Simulated annealing outperforms other approaches on random Ising models. We evaluate our method against 5 benchmarks on 400 randomly generated Ising models that operate on varying sequence sizes. For each model 6 libraries of varying sizes are generated for a total of 2400 experimental conditions. To normalize over the varying conditions, we scale the scores such that the expected score of a library of the desired size generated uniformly at random is 1 unit apart from the maximum possible score of any arbitrary library of the desired size. The scores are then shifted such that SAPS achieves a score of 0, which is indicated by the light gray line in the figures. We do this because the variability of the optimums between different Ising models is much larger than the difference between the approaches, making it difficult to see that SAPS outperforms the other methods in most instances. A) gives the distribution of normalized scores for the approaches we benchmark against using a box plot in conjunction with a violin plot. B) gives the mean of the normalized score for each approach as a function of library size.

#### Theorem 1.

*Let ϵ be any strictly positive constant that is at most 1. Let f*(.) *be expressed as some multi-output boolean circuit. Then unless P* = *NP, there exists no algorithm running in time polynomial to the size of f*(.) *that can, upon receiving f*(.) *as input and L*_*s*_, *L*_p_, *n*_s_, *and n*_p_ *as parameters, find a factorizable library with factors* 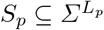 *and* 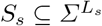 *of sizes n*_p_ *and n*_s_ *respectively such that*:

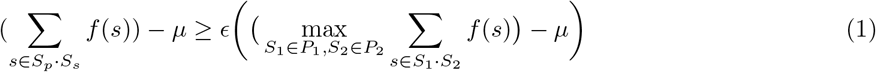

*Where* 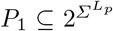 *are all the subsets of* 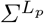 *of size n*_p_ *and let* 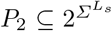 *are all the subsets of* 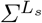 *of size n*_s_, *and* 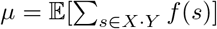, *where X and Y are uniformly random over P*_1_ *and P*_2_.

### 2.3 Stochastically Annealed Product Spaces (SAPS) generates factorizable libraries

We introduce Stochastically Annealed Product Spaces (SAPS) for designing segment libraries that when combinatorially joined optimize a provided objective for the resulting factorizable library. Since producing an efficient factorizable library is intractable, we rely on heuristic methods that employ Gibbs sampling and simulated annealing.

We initially start with randomly generated prefix and suffix libraries with sizes that are chosen to achieve a factorizable library diversity goal. We then perform a Gibbs sampling sweep over all positions in all sequences in both libraries. At each position, we generate all possible substitutions at that position and stochastically accept one of them such that the probability of selecting that update is proportional to the exponential of the score of the concatenated library resulting from that update divided by a temperature parameter. The temperature parameter is then lowered over time according to some schedule so that the stationary distribution approaches an indicator function over the optimal library. This process is illustrated in Figure 1B.

### 2.4 The reverse kernel trick allows for efficient library score evaluation

A major bottleneck of the simulated annealing approach is evaluating the total score of the factorizable library. Modifying a single sequence in the prefix or suffix libraries can affect the scores of 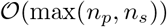 sequences in the factorizable library, so potentially that many reevaluations of *f*(.) would be needed.

Suppose we find a pair of functions 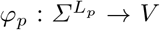 and 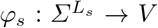 for some inner product space *V* such that *f*(*x* · *y*) = ⟨*φ*_p_(*x*), *φ*_s_(*y*)⟩. Then by the distributive property of inner products:

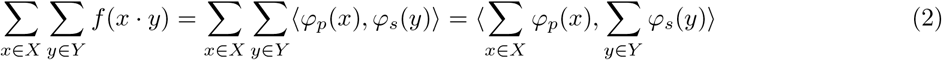

So as long as we keep track of the rLunning sums Σ_*x∈X*_ *φ*_p_(*x*) and Σ_*y∈Y*_ *φ*_s_(*y*) as we update our prefix and suffix libraries, the total score Σ_*x∈X*_ Σ_*y∈Y*_ *f*(*x·y*) can be evaluated by evaluating a single inner product. We will refer to *φ*_p_(.) and *φ*_s_(.) as prefix and suffix feature maps respectively, and we will refer to this optimization as the *reverse kernel trick* in reference to the kernel trick, since in the kernel trick the optimization comes from expressing 〈*φ*(*x*), *φ*(*y*)〉 as a kernel function *k*(*x, y*) for some feature map *φ*(.).

Fortunately, prefix and suffix feature maps that map to finite dimensional Euclidean spaces can be found for any function *f*(.). Theorem 2 shows that the loss of accuracy of computing *f*(.) using the dot product of prefix and suffix feature maps is bounded by the size of the Euclidean spaces employed for *V*.

#### Theorem 2.

*Let* 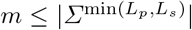 *be a positive integer. Then for any positive integer* 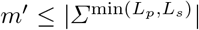, *it is possible to find for every* 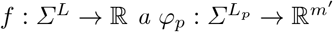 *and* 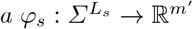 *such that*

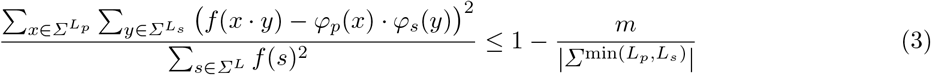

*If and only if m*^′^ ≥ *m.*

Although Theorem 2 does guarantee the existence of feature maps *φ*_p_ and *φ*_s_, it also implies that the dimension of those feature spaces can get very large for certain functions. The reverse kernel trick is unhelpful if the dimension becomes so large that adding vectors and evaluating dot products is inefficient. It is therefore essential to find feature maps with codomains where sums and inner products can be efficiently evaluated.

A special case where features spaces are small is when our scoring function can be described with an Ising model, or more generally a Potts model. Since the only interactions modelled are between pairs of sequence positions, an encoding of size 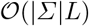 will suffice express all the interactions. Additional details can be found in Appendix B.1.

More generally, we will rely on deep learning to produce these feature maps. We can parametrize *φ*_p_(.) and *φ*_s_(.) with a pair of neural networks, and we can make a *f*(.) predictor by taking the dot product of the outputs of these networks. We can then train this predictor using standard deep learning methodology. Specific details on the architecture and training can be found in Appendix B.3 and B.4.

### 2.5 Sequences of different lengths can be represented using padding

To allow for sequences of differing lengths in the factorizable library, we introduce a padding character in *Σ*. We avoid malformed sequences by pre-specifying which positions in which sequences are padding characters, which ensures that the sequences will always be well formed. A desirable outcome of this approach is that the factorizable library will contain a diversity of sequence lengths. If sequences are instead sampled without prespecified lengths, for instance by allowing the padding characters to be proposed and assigning libraries with malformed strings to have a score of −∞, the tendency will be to sample longer sequences since the number of longer sequences vastly outnumber the number of shorter sequences.

If sequences are fixed length duplicates can be eliminated if we enforce that each sequence in the prefix library is unique and each sequence in the suffix library is unique. This is insufficient if sequences can vary in length. We may for instance propose a prefix library that contains “AC” and “*A”, and a suffix library that contains “D*” and “CD”, where “*” is the padding symbol. Concatenating the two libraries then generates “ACD” twice.

Note that there is no sequence we can remove from the prefix or suffix library without reducing the number of unique sequences in the concatenated library, so it is unclear whether preventing such proposals is desirable. Therefore, we choose to ignore this case and treat the differently padded duplicates in the concatenated library as distinct sequences for library generation purposes. The impact of such duplicates is low: if *Δ* is the difference between the longest and the shortest sequence in the prefix or suffix library, then the number of truly unique sequences in the library can drop by no more than a factor of *Δ* since each sequence can be shifted at most *Δ* times.

### 2.6 Sequence diversity can be explicitly enforced with an entropy term

Beyond attaining efficiency, we would also like a factorizable library to explore a diverse range of sequences. This is partially achieved through the sheer number of unique sequences that factorizable libraries can contain, but this does not necessarily preclude excessive exploitation of certain parts of the sequence space. For example, given a seed peptide of length 20 it is possible to generate a library of size 10^9^ consisting of nothing but mutants that have mutated at most 5 residues away from the initial seed sequence.

To ensure library diversity we add an entropy term to our optimization objective. Let *S*_p_[*i*] ⊆ *S*_p_ be the subset of the prefix library of sequences of length *i*, and let *S*_s_[*i*] ⊆ *S*_s_ be the subset of the suffix library of sequences of length *i*. Let 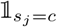 be 1 if *s*_j_ = *c* and 0 otherwise, where *s*_j_ denotes the *j*th letter of *s*. The entropy objective *H* can then be given by the following formula, where for simplicity we define 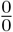 and 0 * *ln*(0) to both evaluate to 0 for the purposes of this formula.

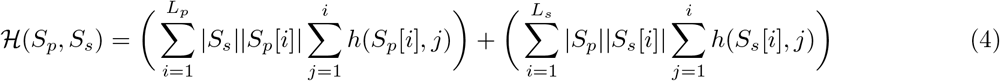

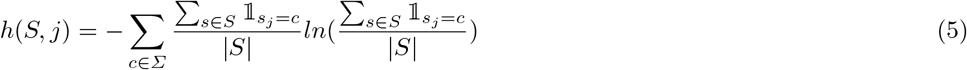

If the prefix and suffix libraries only contains sequences of length *L*_p_ and *L*_s_ respectively, this formula simplifies to the following more interpretable formula:

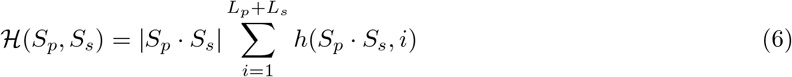

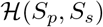 can be thought of as roughly being the number of bits needed to write down every sequence in the factorizable library using an optimal encoding multiplied by *ln*(2). The encoding differs depending on the position along the sequence and the lengths of the prefix and suffix used to generate the sequence it is encoding. Intuitively, the more bits we need to write something down the more diverse it is. For example, this term will incur a penalty if the library consists of a set of mutants that are all close to some seed sequence as described earlier.

#### Theorem 3.

*Let d* ∈ *Σ*^L^, *and let m < L. Let S*_p_ · *S*_s_ ⊆ *Σ*^L^ *be a library where every sequence can be obtained through at most m substitutions of d. Then* 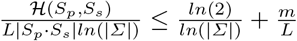.

We remark that *L|S*_p_ · *S*_s_|*ln*(|*Σ*|) can roughly be thought of as the optimal value for 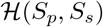, so the above statement characterizes how far the value is from being optimal.

### 2.7 The parameter λ trades off efficiency vs. diversity in library design

We introduce the hyperparameter *λ* that controls the trade-off between the entropy term and the objective function score term that represents efficiency. Using *λ* we can write down the SAPS objective function:

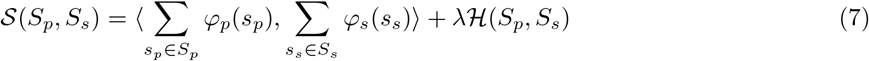

Changes in this quantity induced by changing a single symbol in a single sequence in the prefix library roughly scales with |*S*_s_|. Similarly, changes induced by changing a single symbol in a single sequence in the suffix library roughly scales with |*S*_p_|. An informal derivation can be found in Appendix B.2.

Therefore, to ensure that proposal distributions remain diverse even for large libraries, we divide the score by those quantities before the generating proposal distributions.

## 3 Results

### 3.1 Evaluation of SAPS library design performance on simulated data sets

As a benchmark for the ability of SAPS to produce high scoring factorizable libraries, we first chose a simple design domain. We generate random non-lattice Ising models where coupling energies between any two spins at any two positions are drawn uniformly at random from {−1, 0, 1}, and where the spins at each position has an independent energy also drawn uniformly at random from {−1, 0, 1} (see Appendix C.1 for additional details). The Ising models we generate operate on sequences of length 14, 16, 18, and 20, and we generate 100 models for each length. From a biological perspective, this can be viewed as being analogous to the energy of a peptide threaded through some designed structure under a 2 residue Hydrophobic-Polar scheme evaluated with something akin to a pairwise distance based potential.

We then generate factorizable libraries of sizes 400, 1600, 3600, 6400, 10000, and 14400 for each model that optimize for highest average energy using our proposed approach, where the length of the prefixes and suffixes are exactly half the length of the total sequence. We drop the entropy term for evaluating these factorizable libraries to focus on how the optimization performs.

We benchmarked SAPS against five other approaches, which we sketch out here. Additional details may be found in Appendix C.2. The first approach is the *greedy approach*, where instead of stochastically sampling changes to the libraries we deterministically pick the optimal change, and iterate until convergence. The second approach is to use an *expectation heuristic*, where we determine the average value of a prefix or suffix sequence by averaging over all sequences with that prefix or suffix and then select the top prefixes and suffixes. The third approach is to use a *max heuristic*, where we take the prefixes and suffixes of the top scoring sequences. The fourth and fifth approach are to take the proposals generated by the second and third approaches respectively as a seed proposal, and then apply a *greedy refinement* to it using the greedy approach. Save for the greedy approach, all the approaches we benchmark against would produce the optimal result if there were no couplings between positions in the prefix or suffix.

We find SAPS tends to outperform all other methods. Out of 2400 trials over varying conditions, SAPS achieves the best scoring sequence 2099 times (87%). The greedy approach was able to outperform SAPS 1 of 2400 times, the expectation heuristic 199 of 2400 times (231 times after applying greedy refinement), and the max heuristic 30 of 2400 times (76 times after applying greedy refinement). The scores achieved by different methods in relation to SAPS are given in Figure 6A. When the library size occupies a significant fraction of the sequence space (around 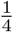 to 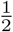), we find that the expectation heuristic tends to outperform SAPS (Figure 6B). This is likely due to how the prefix heuristic becomes a better approximation to the true utility when the suffix library covers a significant chunk of the suffix space, and vice versa. In practice there is rarely a reason to design a library in this regime. If we are allowed such library sizes it would be more sensible to simply use the entire sequence space as a library.

### 3.2 Inner products of small feature vectors produce CDR-H3 enrichment predictions that are comparable to state-of-the-art

We next sought to show that we can learn prefix and suffix feature maps that map to small spaces and can successfully predict affinity to biological targets. We use high throughput sequencing data of three rounds of affinity selection of a random synthetic antibody library against multiple specific targets including the antibodies Ranibizumab, Omalizumab, Trastuzumab, and Etanercept. We also select against a Baculovirus extract (BV), which contains a mixture of viral DNA, proteins, and lipids commonly used to assay polyspecificity of antibody therapeutics in late stage pre-clinical development of a small number of candidates [7]. After each round of selection, antibodies expressed via phage display are isolated and sequenced, hence providing per-round read counts for unique CDR-H3 sequences, which we use to generate training and held-out test sets. For comparison to experimentally generated random synthetic antibody libraries, we collected high throughput sequencing data using the same random synthetic antibody library panned against no target for a single round (FW_kappa), which is the also the same seed library used for previously published phage panning experiments [11]. The majority of sequenced CDR-H3s (~ 99%) in this library range in length from 8 to 20 residues, so we filter out sequences outside this range. We use the *log*_10_-enrichment from round 2 (R2) to round 3 (R3) (*log*_10_(R3/R2)) as a measure of affinity and regression label for this sequence domained prediction task. *log*_10_(R3/R2) enrichment was found to have a better signal-to-noise ratio when compared to the inclusion of round 1 (R1) reads and previous work has shown that this measure correlates well with ground-truth CDR-H3 affinity measured by individual binding assays, such as enzyme-linked immunosorbent assay (ELISA) [11].

We use deep learning to find prefix and suffix feature maps that map to low dimensional feature spaces (see Appendix B.3 and B.4). Specifically, we use a deep convolutional neural network with residual connections (i.e. a ResNet) to map prefixes and suffixes to 16-dimensional vectors. The inner product between a prefix and suffix feature vector then gives the predicted *log*_10_(R3/R2) enrichment. We will refer to this entire pipeline as a reverse kernel model.

It is possible for the reverse kernel model to appear to perform well even if it fails to capture the non-linear interactions between the prefix and the suffix positions of a sequence if those interactions are sufficiently negligible. To control for this, we train a pair of ResNets, where one predicts a score on the prefix and one predicts a score on the suffix. The scores are then added to produce the overall prediction. We will refer to this pipeline as the independent model. We use this as a baseline to evaluate how well the reverse kernel model captures the non-linear interactions.

Finally we also trained a ResNet to output the scores without any intermediate steps, which we will refer to as the unrestricted model. We will rely on this model to evaluate our designed libraries, since we expect it to produce the most accurate approximation of the true underlying sequence landscape due to its ability to share information between the prefix and the suffix in any of its layers, allowing it to capture a more nuanced perspective of that landscape.

We compare the performance of our models by computing the Pearson r correlation between the predicted and observed *log*_10_(R3/R2) enrichment on held out validation sets. The results are presented in Figure 3, where we see that generally the unrestricted model does indeed perform the best (with the one exception of Ranibizumab), suggesting that it does provides the best approximation of the sequence landscape. We also see that the reverse kernel model outperforms the independent model, which demonstrates that it is able to combine prefix and suffix properties in a non-linear way to produce better generalizations. Our reverse kernel models attain Pearson r values of 0.83, 0.64, 0.63, 0.65 and 0.87 on validation sets for Ranibizumab, Trastuzumab, Omalizumab, Etanercept, and BV respectively, which is comparable to the values that were reported in prior work, which were 0.79, 0.65, and 0.64 for Ranibizumab, Trastuzumab, and Etanercept respectively [11].

**Fig. 3.**
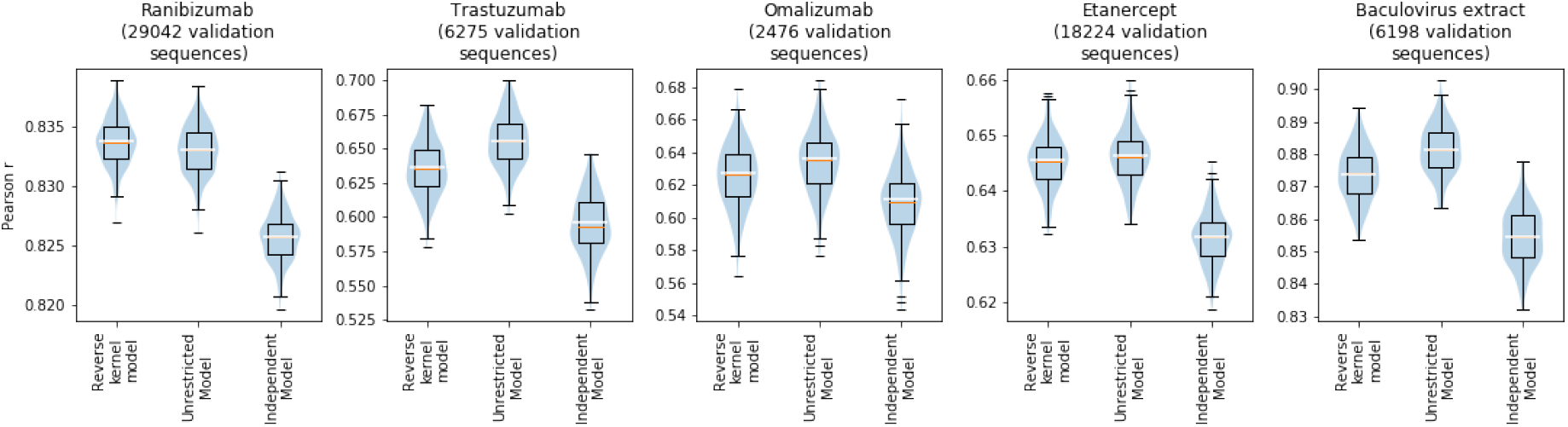
Model performance. We compare the performance of our reverse kernel models with that of our unrestricted models and independent models on validation data. The Pearson r values on the validation set are indicated with a white bar in the above plots, and we use a box plot in conjunction with a violin plot to show the uncertainty as measured using 250 bootstrap samples of the validation data set.

### 3.3 SAPS produces diverse factorizable libraries with optimized affinity for specific targets

To generate the factorizable libraries, we take the reverse kernel models we trained and performed the SAPS procedure for 500 sweeps, decreasing the temperature by a factor of 1.1 every 5 rounds. For the purposes of SAPS, the values output by the reverse kernel models were divided by their standard deviation, which was estimated with 100000 randomly generated sequences following the same length distribution as the factorizable library, with residues generated uniformly and independently.

We generated pairs of prefix and suffix libraries of each target that each contain 35,000 sequences. When combined, they produce factorizable libraries containing over 10^9^ designed sequences that are optimized for binding to Ranibizumab, Omalizumab, Trastuzumab, and Etanercept (see Appendix D.2 for additional details). The diversity hyperparameter *λ* used for designing these libraries was chosen by observing its effects on smaller libraries (see Appendix D.3). Generally, we recommend using hyperparameters between 0.1 and 0.3 depending on whether a user’s priority is exploration or exploitation of the sequence space.

We find that our libraries are highly diverse in comparison to FW_kappa, the experimentally randomized synthetic antibody library used in the previously described phage display and affinity selection experiments. The expected Levenshtein distances between a pair of randomly selected sequences of length 12 within our libraries are around 2 edits further than the expected Levenshtein distances between a pair of randomly selected sequences of length 12 within FW_kappa, and are around only one edit closer than the expected Levenshtein distances between a pair of uniformly random sequences of length 12. While FW kappa serves as an approximation of the diversity of a random library, we note that it may contain a biased sequence distribution due to the round of no-target panning and other experimental constraints prior to high-throughput sequencing, similar to other proposed antibody CDR-H3 libraries [9]. The full distributions are presented in Figure 4A. Next, we show that diversity is similar in the prefix and suffix region for each factorizable library and is not derived from one side of the CDR-H3 sequence alone in Appendix E.2.

**Fig. 4.**
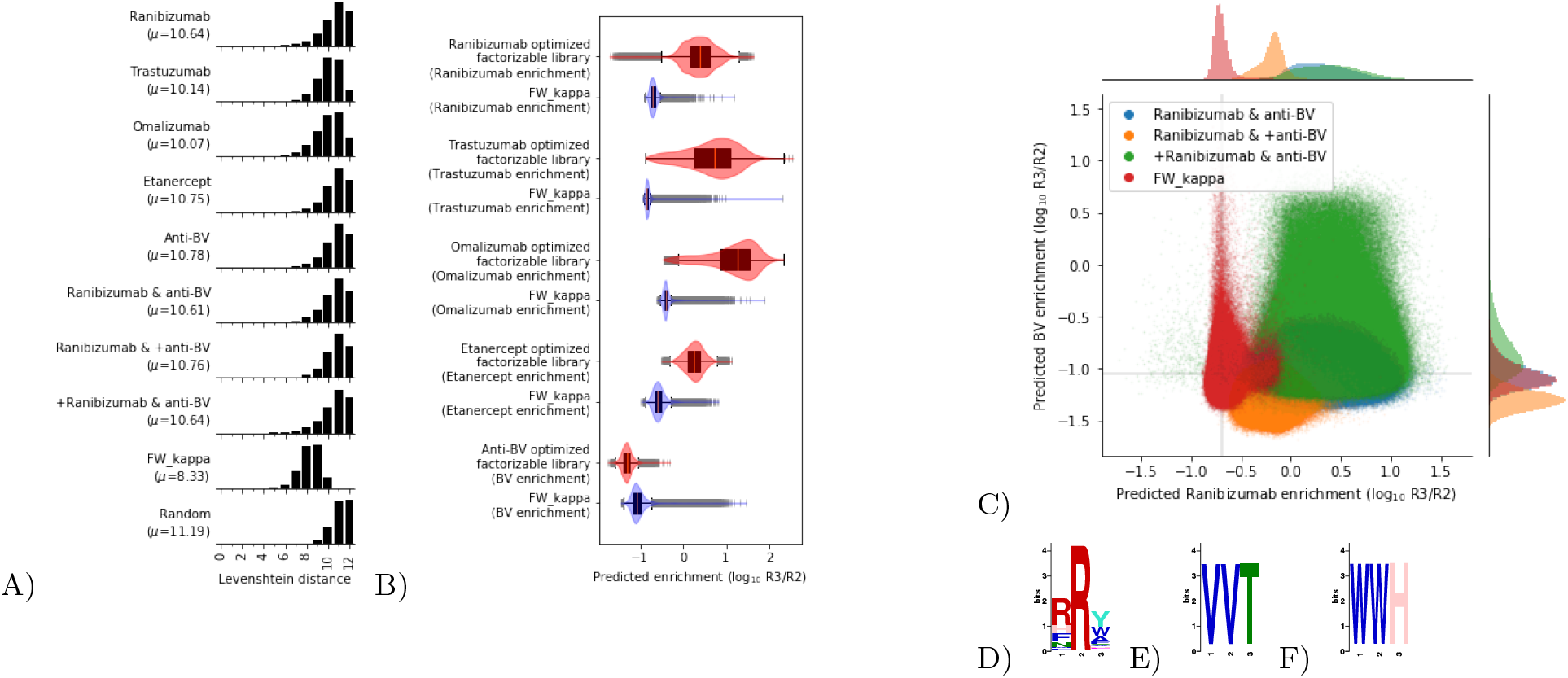
Library validation We compare the diversity of each library that was generated using SAPS and affinity selection data to FW kappa. A) Histograms are shown of the pairwise Levenshtein distance between pairs of length 12 sequences in each library. Also included is the histogram obtained when pairs of sequences are uniformly random. The mean distance, *μ* is reported. B) We compare the sequence optimality of generated libraries against FW kappa by scoring the sequences of FW_kappa and 1000000 uniformly random samples from the designed libraries with the corresponding unrestricted model. Score distributions are reported as boxplots laid over violin plots for FW kappa (blue) against each designed library (red). C) Joint plot shows predicted enrichment of FW kappa (red), equally weighted libraries (blue), Ranibizumab weighted libraries (green) and anti-BV weighted libraries (yellow) by the Ranibizumab unrestricted model on the x-axis and the BV unrestricted model on the y-axis. Panels D-F show SeqLogos for enriched nonspecific motifs in the Ranibizumab weighted library over the anti-BV weighted library discovered by STREME. D) `RRY` motif (p-value=4.8 * 10^*−*2315^). E) `VVT` motif (p-value=4.2 * 10^*−*38^). F) `WWH` motif (p-value=9.8 * 10^*−*21^).

We also find that our libraries score significantly better on enrichment for the targets they were optimized for in comparison to FW_kappa. The target specific enrichments were estimated by running the corresponding unrestricted model. The distributions of the enrichment scores are presented in Figure 4B. Together, these results suggest that SAPS is able to generate factorizable libraries that are both diverse and efficient. Details on pairwise distance computation and model scoring are provided in Appendix E.1.

### 3.4 Flexible SAPS parameters allow for design of limited polyspecificity factorizable libraries

Next, we show that SAPS can produce factorizable libraries with specified diversity and sequence optimality constraints. In this task, we focus on designing a library with limited polyspecificity using the aforementioned BV target. Generally, if an an antibody binds a target like BV, it is likely a polyspecific sequence that will bind many different targets in the body leading to diminished efficacy by fast immune clearance or even clinically dangerous off-target effects [5].

We used SAPS to design a factorizable library with *low* affinity for BV, or “anti-BV”. This is done by negating the output of the reverse kernel model, and is intended to reduce the number of polyspecific members included in the libraries. We show that this library has high diversity but lower affinity for BV than the FW_kappa library, indicating a better polyspecificity profile in Figure 4A and B. Further, we conduct basic motif enrichment analysis and show that hypothesized nonspecific motifs, as theorized by [9], are less prominent in the designed factorizable library (see Appendix E.3).

Next, we used SAPS to design factorizable libraries that were optimized for both increased affinity to Ranibizumab *and* for decreased affinity to BV. To achieve this, we use a weighted sum of the Ranibizumab reverse kernel model output and the negated BV reverse kernel model output to score sequences for our optimization objective. We generate three factorizable libraries, one where both outputs are unscaled (“Ranibizumab & anti-BV” or “equally weighted”), one where Ranibizumab is scaled by 0.1 while anti-BV is unscaled (“Ranibizumab & +anti-BV” or “anti-BV weighted”), and one where anti-BV is scaled by 0.1 and Ranibizumab is unscaled (“+Ranibizumab & anti-BV” or “Ranibizumab weighted”). We evaluate efficiency by scoring each library with unrestricted models predicting Ranibizumab affinity and BV affinity, showing that the libraries designed have the intended score distribution (Figure 4C). Further, we conducted motif enrichment in the Ranibizumab weighted library (+Ranibizumab & anti-BV) over the anti-BV weighted (Ranibizumab & +anti-BV) library using STREME (see Appendix E.3 for details) and observe significant enrichment of known nonspecific motifs such as valine (VV), tryptophan (WW) and arginine (RR) pair enrichment. These results are presented in Figure 4D-F, and demonstrate that SAPS is flexible for highly specific design parameters. This also illustrates a functional use-case for rationally designed factorizable libraries, as it is common in antibody discovery to spend significant resources on both finding a sequence with optimized affinity for a target while rejecting sequences with high polyspecificity [14].

## 4 Discussion

We present SAPS, a computational method to guide the synthesis of *factorizable libraries*, a novel synthesis strategy that enables the rational design of highly diverse libraries with optimized properties at moderate cost. As a result of their skewed focus on exploration and exploitation of the sequence space, respectively, random libraries and the individual design of every sequence in a library are not ideal for the discovery of novel therapeutics, especially for difficult targets. Further, it is currently not feasible to create libraries with 10^9^ rationally designed members by direct synthesis. With rationally designed *factorizable libraries*, smaller segment libraries are synthesized at low-cost and combined to produce a full-length library that is orders of magnitude larger. By guiding the design of the segment libraries, factorizable giga-libraries can contain a higher proportion of optimized sequences for use in therapeutic selection experiments, increasing the probablity of discovering novel therapeutics targeting difficult biological and disease targets. We demonstrate SAPS by designing factorizable antibody CDR-H3 sequence libraries against various targets. We note that SAPS designed factorizable libraries can be used for any discovery task that can benefit from the direct synthesis of diverse and functionally efficient sequencing library design tasks such as TCR libraries [19, 1], AAV capsid libraries [4, 10, 18], DNA/RNA libraries such as aptamers [13, 8], as a few examples.

We also note that independently designed factorizable libraries can be synthesized, ligated, and subsequently mixed to form a single *integrated factorizable library*, that integrates the objective functions of each of the underlying factorizable libraries. This method allows dependencies between segments to be captured in each component factorizable library.

We have shown that reverse kernel models can reliably recapitulate sequence-function relationships as measured by experimental affinity selection. We demonstrate that these models can be used as a scoring function for SAPS to generate factorizable libraries that, upon combination, contain 10^9^ members. We show that these factorizable libraries explore the sequence space by computing the pairwise edit distances between sequences in these giga-libraries showing their superior diversity when compared to an experimentally generated random synthetic library. By scoring generated sequences using validated unrestricted models, we show that designed factorizable libraries are more efficient than randomized libraries and reflect intended objectives for given tasks and exploit the sequence landscape. Finally, we show that our method flexibly allows for fine-tuning of design parameters such as the overall edit distance between sequences and trade-offs between multiple desired sequence properties.

## 5 Data access

Code and training data are available at https://github.com/gifford-lab/FactorizableLibrary.

## 6 Competing interest statement

Geraldine Horny, Christine Banholzer, and Stefan Ewert are employees of Novartis. The remaining authors declare no competing interests.

## 7 Acknowledgements

This work was funded by NIH Grant R01 CA218094 and a gift from Schmidt Futures to D.K.G, and the experimental work was funded by Novartis.

## Appendices

### A Proofs of statements

#### A.1 Proof of Theorem 1

We will prove the contraposition and show that P = NP is implied. If the algorithm described in the statement exists, then there exists an algorithm *A* that can, upon being given *f* : *Σ*^2L^ → {−1, 0, 1} as a 2 bit output boolean circuit and *n* ≤ |*Σ*^L^|, return within time polynomial in *f*(.) a pair of sets *S*_p_ ∈ *Σ*^L^ and *S*_s_ ∈ *Σ*^L^ of size *n* such that:

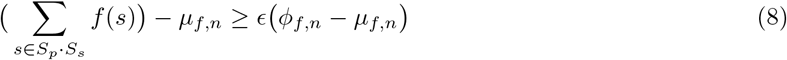

Where *μ*_f,n_ denotes the expected score of a random solution and *ϕ*_f,n_ denotes the optimal solution. Let 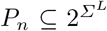 be all subsets of *Σ*^L^ of size *n*, and let *X* and *Y* be drawn uniformly at random from *P*_n_. Then formally:

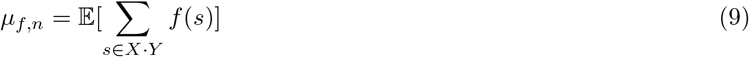

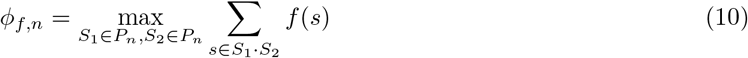

Let *G* = (*V, E*) be some graph that we wish to approximate the max clique of. Let *L* = ⌈*log*_|Σ|_(|*V*|^3^)⌉, so |*V*|^3^ ≤ |*Σ*^L^|. We then pick a subset of *V*^′^ ⊆ *Σ*^L^ to represent the vertices *V*. We can then construct a function *f* : *Σ*^2L^ → {−1, 0, 1}:

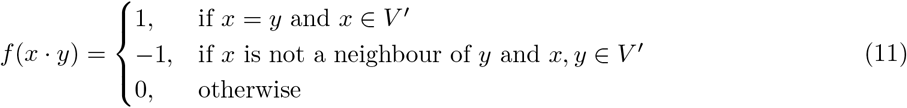

We can express *f* as circuit that is polynomial in the size of *G* by constructing it to match sequences with prefixes and suffixes in *V*^′^ and ignoring all other sequences and outputting 0. It is also clear that it can be constructed in time polynomial to |*V*^′^|. We then feed *f* to algorithm *A* along with *n* = |*V*|. Let *X* be the score of the solution found by *A*.

##### Lemma 1.

*Suppose n* ≤ |*V*^′^|. *Then μ*_f,n_ ≥ |*V*^′^|^−2^.

*Proof.* First we note that for any *S*_1_ ⊆ *Σ*^L^ and *S*_2_ ⊆ *Σ*^L^, we have 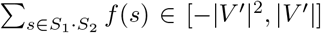. This is because at most |*V*^′^| sequences evaluate to 1 and at most |*V*^′^|^2^ sequences evaluate to −1.

Next, consider the probability of drawing a random subset of *Σ*^L^ that has size *n* that contains at least one sequence in *V*^′^. The probability of this is upper bounded by the expected number of sequences in *V*^′^ in such a subset, which is 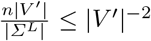 by construction. Therefore, the probability of randomly drawing two subsets *S*_1_, *S*_2_ ⊆ *Σ*^L^ that have size *n* such that 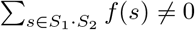 is at most |*V*^′^|^−4^.

Therefore, 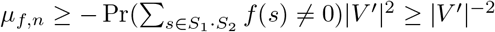, where *S*_1_ and *S*_2_ are uniformly random subsets of *Σ*^L^ of size *n*, as desired.

##### Lemma 2.

*Let n* = |*V*^′^|. *Then ϕ*_f,n_ *is equal to the size of the max clique in G*.

*Proof.* The only way a we can contribute positively to the score is if an element *v* belonging to *V*^′^ is contained in both libraries. However, this contribution is negated if there is some sequence in *V*^′^ contained in either library that is not a neighbour of *v*. Therefore, the best score can be no larger than the size of the max clique in *G*. Conversely, we can attain the size of the max clique as a score if we include the elements in *V*^′^ that form the clique in both libraries, and make every other sequence elements outside of *V*^′^.

Then we have the following:

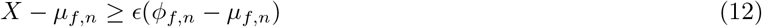

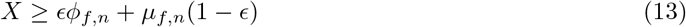

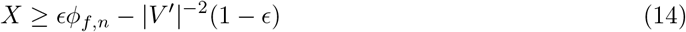

For sufficiently large graphs, 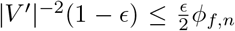 since *ϕ*_f,n_ ≥ 0 (we can include a single sequence of |*V*^′^| in both libraries and make sure every other sequence is not in |*V*^′^|). Therefore:

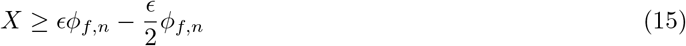

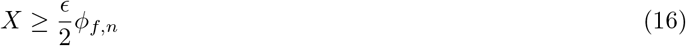

Therefore, *X* is at least a 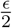 fraction of the size of the max clique. Thus, we have constructed a polynomial time procedure for attaining a 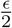 approximation of the size of the maximal clique. Since any constant factor approximation of the size of the maximum clique is known to be NP-hard [20], we have *P* = *NP* as desired.

We remark that the result from [20] in fact implies a stronger inapproximability result, but we omit this to simplify the statement.

#### A.2 Proof of Theorem 2

Let *A*_x,i_ denote the *i*th entry in the vector *φ*_p_(*x*). Let *B*_y,i_ denote the *i*th entry in the vector *φ*_s_(*y*). Let *F*_x,y_ denote *f*(*x · y*). If we assign a total ordering to elements of the prefix and suffix spaces and treat elements of those spaces as the numbers denoting their ordering, then we can view *A*, *B*, and *F* as matrices. We can then write Equation 3 as the following:

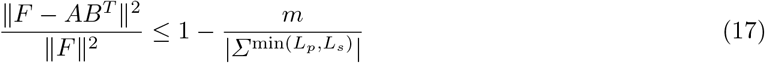

Where ||.|| denotes the Frobenius norm of a matrix. By definition, *AB*^*T*^ is a rank *m*^′^ matrix. Let *USV* = *F* be the singular value decomposition of *F*. By the Matrix Approximation Lemma, ||*F − AB*^*T*^|| is minimized when 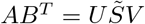, where 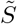 is *S* where only the largest *m*^′^ values along the diagonal are not zero. Therefore:

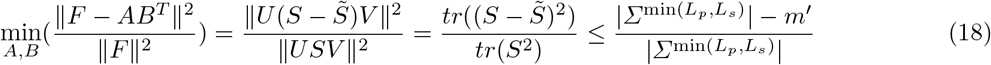

Therefore, Equation 17 holds for all *F* if *m*^′^ ≥ *m*. If *m*^′^ *< m*, then we note that the last inequality in Equation 18 holds with equality if *S* is a matrix with zero entries everywhere except along the main diagonal, which implies that there is some *F* for which Equation 17 does not hold for any *A* and *B*. Theorem 2 follows.

#### A.3 Proof of Theorem 3

Let *p*_i,c_ denote the fraction of sequences with character *c* at position *i*. We can then write 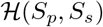 as the following:

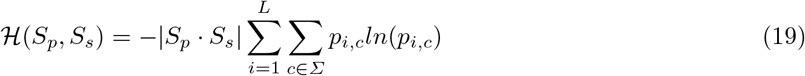

We also have the following relations:

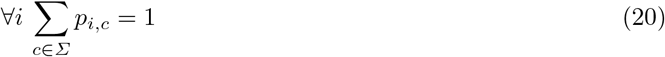

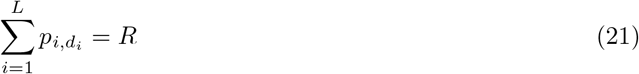

Where *R* is some constant that is at least *L − m* and *d*_*i*_ denotes the *i*th character of *d*. The first relation follows since the proportions must add up to 1, and the second relation follows since all sequences can be obtained with *m* substitutions of *d*.

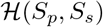 is upper bounded by the maximum value 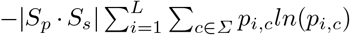 can take on subject to the constraints imposed by Equations 20 and 21. The optimization problem can be solved via Lagrange multipliers, which yields the following critical point:

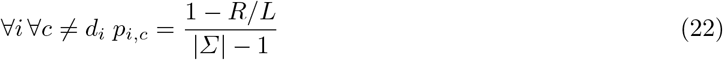

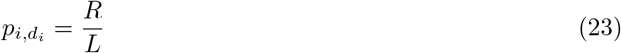

Since this is the only critical point, and since it is straightforward to find *p*_i,c_ under the constraints such that the objective evaluates to a lower value (set 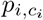 to 1 *R/L* + *ϵ* for all *i* where *c*_i_ ≠ *d*_i_ is some arbitrarily chosen symbol and where *ϵ* approaches arbitrarily close to 0), this must be a global maximum. Therefore:

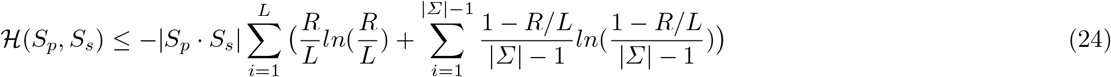

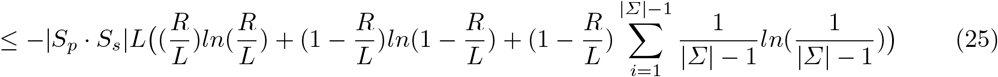

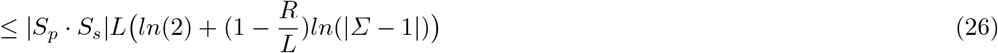

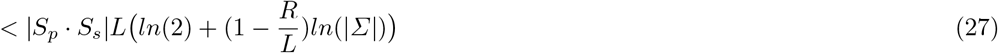

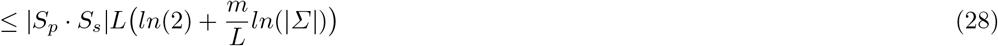

The final inequality follows from *R ≥ L − m*. Dividing the overall inequality by *L|S*_p_ · *S*_s_|*ln*(|*Σ*|) yields the desired result.

### B Additional details for methods

#### B.1 Small feature spaces for Potts models

If *f*(*x*) gives the energy of *x ∈ Σ*^L^ and can be described by a Potts model, then a pair of feature maps 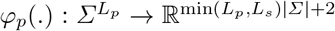 and 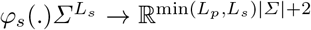 can be found. First, we express sequences *x ∈ Σ*^L^ as a tensor where *x*_i,c_ = 1 if *x*_i_ = *c* and 0 otherwise. Then there must exist some rank 4 tensor *A* such that the following holds:

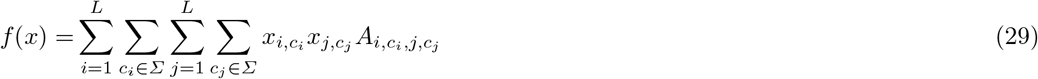

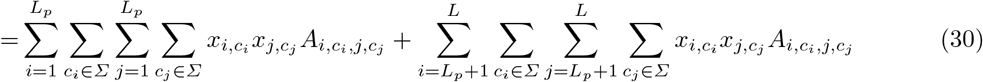

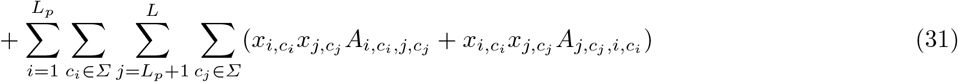

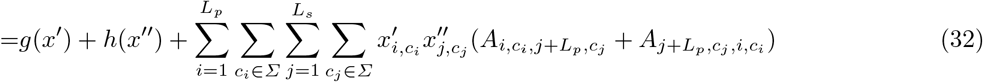

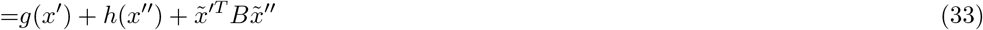

Where *x*^′^ is the matrix representing the prefix of *x* and *x*^′′^ is the matrix representing the suffix of *x*. 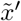 and 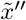 are the flattened vectors of *x*^′^ and *x*^′′^, and *B* is some *L*_p_|*Σ*|-by-*L*_s_|*Σ*| matrix obtained by reshaping and summing the appropriate subtensors of *A*.

Without loss of generality, let *L*_p_ ≤ *L*_s_. Then we can simply let 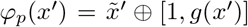 and 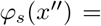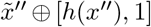, where ⊕ denotes vector concatenation. Therefore both prefix and suffix feature maps map to 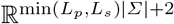, and *φ*_p_(*x*^′^) · *φ*_s_(*x*^′′^) = *f*(*x*) as desired.

#### B.2 The objective function scales with library size

Suppose we propose to change a sequence *x* to *x*^′^ in the prefix library, where *x* and *x*^′^ differ by only a single residue at position *i*. Let *l* be the length of *x* and *x*^′^. Let 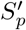 denote the updated prefix library. The difference in objective between the initial library and the proposed library is then:

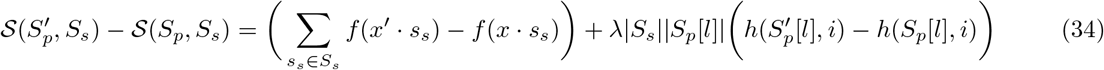

The first term has a sum that contains |*S*_s_| terms. If our proposal is an improvement to most sequences it affects, then the size of the sum roughly scales with |*S*_s_|. For the second term, if the library is sufficiently large we can estimate the change in entropy with the derivative:

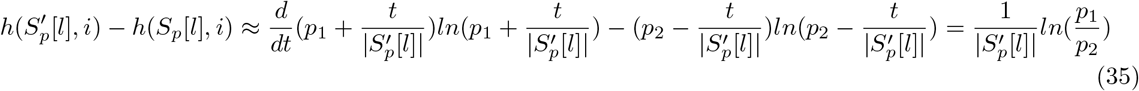

Where *p*_1_ is the fraction of sequences in *S*_p_[*l*] that has *x*_i_ at position *i* and *p*_2_ is the fraction of sequences in *S*_p_[*l*] that has 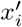 at position *i*. Thus the second term in equation 34 also scales with |*S*_s_|.

So the overall difference in score also scales with |*S*_s_|. Similarly, we can see that making a change to a sequence in the suffix library induces a change that scales with |*S*_p_|. Thus in order to maintain a stochastic phase in the simulated annealing procedure we must normalize the score.

#### B.3 Deep learning architecture

We employ a convolutional neural network with residual connections to map strings to real valued vectors. First, we strings are encoded into a 40-by-*n* array, where *n* is the maximum size of the string that the model can take as input. This is done by first padding out the string to maximum length, and taking each character in the string and mapping them into a 40 dimensional embedding, where the first 2 0 e ntries c onsists o f a one-hot encoding denoting the residue and the last 20 entries is the BLOSUM62 substitution values [22]. The padding character is assigned the zero vector. These vectors then make up the rows of the resulting encoded matrix and is fed into the neural network, where convolutions are run over the second dimension and the first dimension i s treated as a channel dimension.

The model first applies a linear transform to each position that increases the number of channels from 40 to 64, followed by batch normalization. This is then run through 5 residual blocks. Each residual block consists of a pair of 1D convolutions with a kernel of size 3 and batch normalizations, separated by a ReLU layer in between. The input of the block is then added to the output, forming the residual block. Max pooling over adjacent positions is preformed after the last 2 residual blocks. The resulting matrix is then flattened and linearly mapped to a 128 dimensional vector, which is then passed through a ReLU layer, which is then linearly mapped to an output vector.

For this work we make use of three kinds of models. The unrestricted model accepts length 20 sequences and outputs 1 dimensional vectors, which are treated as output scores. The independent model consists of a prefix and a suffix model that accept length 10 sequences and output 1 dimensional vectors, which are added together to give the final output scores. The sequence is first padded to length 20, and then divided into two sequences of length 10 which are fed into the prefix a nd s uffix mod els. The rev erse ker nel mod el operates identically to the independent model, except the models output 16 dimensional vectors. The dot product of the vectors then give the final score.

#### B.4 Training deep learning models

Training datasets were split into 10 equal sets (one set has slightly more or less sequences when the number of sequences is not a multiple of 10). These were used to create 10 training and validation splits, where for each split one of the sets was used for validation and the rest were combined for training. For each split, two randomly intialized models were trained, resulting in an ensemble of 20 models. Final predictions are made by averaging over the outputs of the ensemble. Each model was trained for 100 epochs using ADAM with default PyTorch v1.7 parameters [24]. Model performances were evaluated using the validation set after each epoch, and the model with the highest performance was saved. Models only accept sequences of length 20, so shorter sequences were randomly padded such that each shift occurs with equal probability. This allows the model to learn shift invariance, which is necessary for sequence generation since different shifts may be represented once prefix and suffix libraries are concatenated. Models were created using PyTorch v1.7 and trained on either a single NVIDIA Titan RTX GPU (24GB RAM) or a single GeForce GTX 1080 Ti (11GB RAM).

Held out validation sets were used to evaluate performance in Figure 3. These sets were filtered to exclude any sequences that overlapped the training sets. Sequences were padded equally on both sides during this evaluation, with extra padding applied to the end if the sequence length is odd.

### C Additional details on benchmarking SAPS

#### C.1 Additional details on the problem domain

To test SAPS, we generate energy landscapes over the domain of fixed length binary strings that are defined by non-lattice Ising models. The Hamiltonian takes the following form:

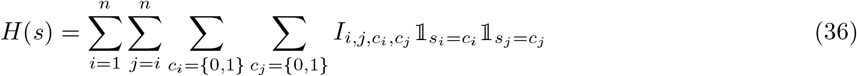

Where *n* is the length of the strings, *s* denotes a string of length *n*, and 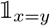 is the indicator function that returns 1 when *x* = *y* and 0 otherwise. To generate a single Ising model, 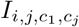 are all independently and uniformly drawn from {0, −1, 1}. 100 such models were generated for domains over sequences of length 14, 16, 18, and 20 for a total of 400 models.

This model permits us to define a a pair of practical feature maps that allows us to use the reverse kernel trick as described in Appendix B.1.

Such a model can be seen as a toy model for protein design. For example, suppose we know the exact desired locations and orientations of the alpha carbons of the protein. That geometry then determines the interactions, which can then be captured with something akin to the model described above.

#### C.2 Alternative optimization approaches to SAPS

The first approach we benchmark against is the greedy approach, where we start with a randomly generated pair of libraries. We then sweep over each bit in each sequence in each library, flipping it if it produces an improvement. We keep sweeping until convergence. Note that this is equivalent to SAPS if the temperature parameter approaches zero.

For the next two approaches, we take each segment library and rank the sequences according to some heuristic. The two use are the expectation and max heuristic. For the expectation heuristic, we take a segment and assign it the mean of all sequences containing that segment. For the max heuristic, we assign to the segment the optimal score of a sequence containing that segment. We then take the top scoring segments from each segment libraries.

Note that in most cases even these heuristics must be approximated since finding the top scoring segments may be itself intractable. However our benchmarking landscapes and functions are small enough such that we can calculate these values exactly. Thus, our benchmarks represent an optimistic view for how well these approaches can perform.

In fact, if there are no interactions between the segments these heuristics will give the optimal solution. Having no interactions means the Hamiltonian of the Ising model would take on the following form:

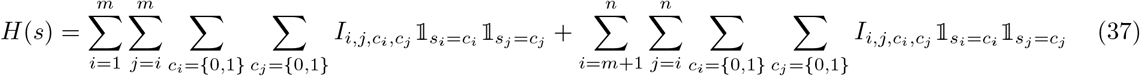

Where the lengths of the strings is *n*, the lengths of the prefixes is *m*, and the lengths of the suffixes is *n* − *m*. We present the proof:

*Proof.* Without loss of generality, suppose we are optimizing for libraries with high scores. Let *X* and *Y* denote the set of prefixes and suffixes respectively. For any string *x* · *y* where *x* ∈ *X* and *y* ∈ *Y* is a suffix, we have *H*(*x* · *y*) = *H*_p_(*x*) + *H*_s_(*y*) for some *H*_p_(.) and *H*_s_(.).

Let *E*_p_(.) denote the expectation heuristic on prefixes and let *M*_p_(.) denote the max heuristic on prefixes. The *E*_s_(.) and *M*_s_(.) denote these heuristics on the suffixes. Let *x* ∈ *X*. We have 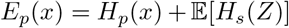 and *M*_p_(*x*) = *H*_p_(*x*) + max_z∈Y_(*H*_s_(*z*)), where *Z* denotes a random suffix distributed uniformly. Note that the second term is independent of *x*, therefore both heuristics would rank prefixes identically, and propose the prefixes with the highest *H*_p_(*x*). The identical argument shows that the heuristics would propose the suffixes with the highest *H*_s_(*y*).

Therefore, it suffices to show that the expectation heuristic provides the optimal solution. Suppose that there exists some pair of libraries that has a higher score than what’s given by the heuristic. Then there must be some segment *x* that is in the better library and not the heuristic library, and some segment *y* that is in the heuristic library but not in the better library such that *E*_p_(*x*) ≤ *E*_p_(*y*) (or *E*_p_(*x*) ≤ *E*_s_(*y*)). Then swapping out *x* for *y* can not decrease the scores of any sequence, since *H*_p_(*x*) ≤ *H*_p_(*y*) (or *H*_s_(*x*) ≤ *H*_s_(*y*)). If we keep swapping we will eventually obtain the library given by the heuristic. Since each swap cannot decrease the score, the heuristic library cannot score worse than the better library, which is a contradiction.

For the final two approaches, we take the greedy approach, but instead of initializing with a random library we initialize with the outputs of the expectation and max heuristics. We refer to this as “greedy refinement” in the main text.

### D Experimental details

#### D.1 Details of phage panning experiments and processing

The experimental single framework library (used in all panning experiments) was constructed as follows: a gene fragment encoding the germline framework combination IGHV3-23 and IGKV1-39 was synthesized in Fab format and cloned into a phagemid vector template. Only CDR-H3s were diversified via trinucleotide synthesis technology. Three of the four publicly available antibody-antibody targets were collected in-house and reported on in [11]. Ranibizumab is a Fab fragment that binds to vascular endothelial growth factor A, Etanercept is a fusion protein that fuses the TNF receptor 2 to the Fc and hinge region of a IgG1 heavy chain, and Trastuzumab.The fourth, Omalizumab, a IgG1k monoclonal antibody that specifically binds to free human immunoglobulin E, and BV was collected in the same fashion as follows: First, the targets were expressed in human IgG1 format. Three rounds of panning are completed in solid-phase mode on 96-well maxisorb plates that are coated with the target with decreasing concentrations over rounds. A negative control panning against no target for one round was conducted and is referred to as FW-kappa in the text. Panned phages are eluted and propogated for high throughput sequencing via MiSeq or HiSeq sequencers. For all experiments, after obtaining sequencing reads, the fixed flanking sequences on the boundary of the variable region were used as a template to BLAST short read alignment (allowing 3 mismatches on each side) and segment out CDR-H3 seqeunces. Datasets were constructed by retaining only sequences that had more than 5 read counts in at least one panning round or had non-zero reads in all rounds to reduce noise.

#### D.2 Generating the factorizable library

We used the phage panning data to train reverse kernel models, which we used to generate segment libraries that combine into libraries of size 10^9^. Each segment library contains 5000 sequences of each length between 4 to 10, for a total of 35000 sequences. The resulting factorizable library should have a length distribution similar to the convolution of a pair of uniform distributions, which is shaped like an isosceles triangle. Sequences in the prefix library were padded exclusively on the left while sequences in the suffix library were padded exclusively on the right before being fed to the reverse kernel models, so the model would not detect gaps in the sequences.

As discussed in Section 2.5, there may be duplicates, so the libraries may not contain exactly 35000^2^ sequences. The number of unique sequences in each library is given in Table 1, along with the *λ* entropy hyperparameter that was used when generating them.

**Table 1.**
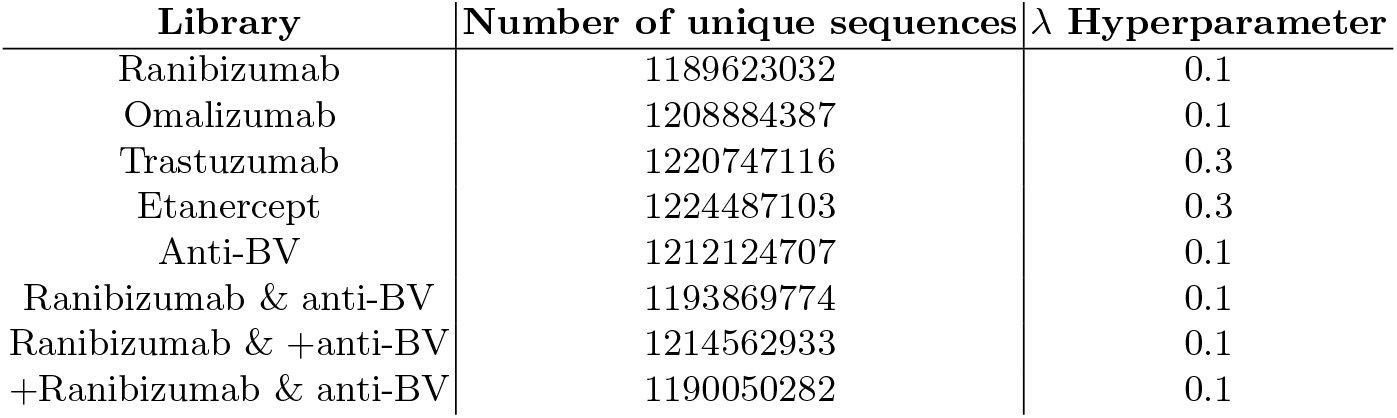
Unique sequences in factorizable libraries and their entropy hyperparameter

| Library | Number of unique sequences | $\lambda$ Hyperparameter |
| --- | --- | --- |
| Ranibizumab | 1189623032 | 0.1 |
| Omalizumab | 1208884387 | 0.1 |
| Trastuzumab | 1220747116 | 0.3 |
| Etanercept | 1224487103 | 0.3 |
| Anti-BV | 1212124707 | 0.1 |
| Ranibizumab & anti-BV | 1193869774 | 0.1 |
| Ranibizumab & +anti-BV | 1214562933 | 0.1 |
| +Ranibizumab & anti-BV | 1190050282 | 0.1 |

#### D.3 λ Hyperparameter tuning

To select *λ* which is a weight on the entropy term in the objective function for model training, we first generate smaller 10^5^ libraries composed of segment libraries that contain 700 segments each (100 segments for each length between 4 to 10 inclusive) at *λ* = [0, 0.001, 0.01, 0.03, 0.1, 0.3, 1]. We then evaluate the pairwise Levenshtein distance between members of this library and score the library with the corresponding unrestricted model. Based on this analysis across targets, we recommend setting this hyperparameter between 0.1 and 0.3 based on the intended library diversity or score distribution. We provide an example of this tuning conducted on Ranibizumab in Fig. 5).

**Fig. 5.**
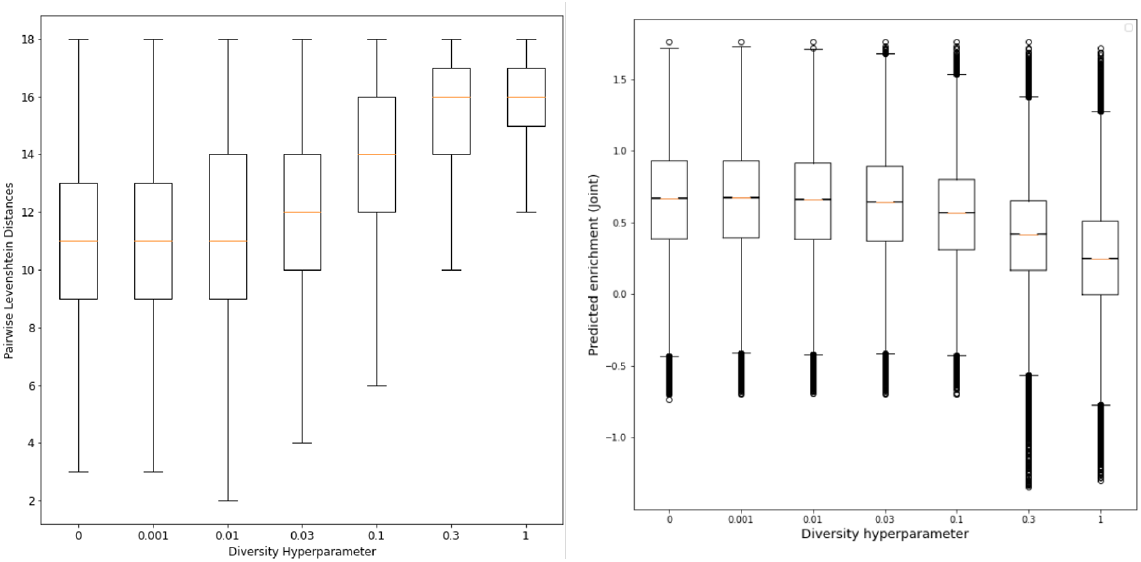
Ranibizumab *λ* hyperparameter tuning

### E Details on Analyses

#### E.1 Details of library validation metrics

For all examples of library validation against FW-kappa, we evaluate diversity by measuring the pairwise Levenshtein distance between 10,000,000 uniformly sampled pairs of length 12 sequences. Briefly, the Levenshtein distance between two strings is the minimum number of single-character edits (substitutions, insertions, or deletions) required to change one string into the other. We compute Levenshtein distance using python-Levenshtein v0.12.2. For gigalibrary scoring, 1,000,000 sequences are uniformly sampled and scored with the corresponding unrestricted model to evaluate the sequence optimality of large generated libraries.

#### E.2 Diversity of prefix and suffix libraries independently

For diversity analysis, we have primarily shown that the diversity of full length SAPS designed CDR-H3 sequences is higher than FW_kappa. To further investigate the source of diversity, here we show that the individual Levenshtein distance distributions of the prefix and suffix libraries are similar across target objectives. Each distribution is calculated from 10000000 uniformly sampled segments over all segment lengths. This indicates that diversity of concatenated libraries is not a result of diversity isolated to the prefix or suffix of the CDR-H3.

**Fig. 6.**
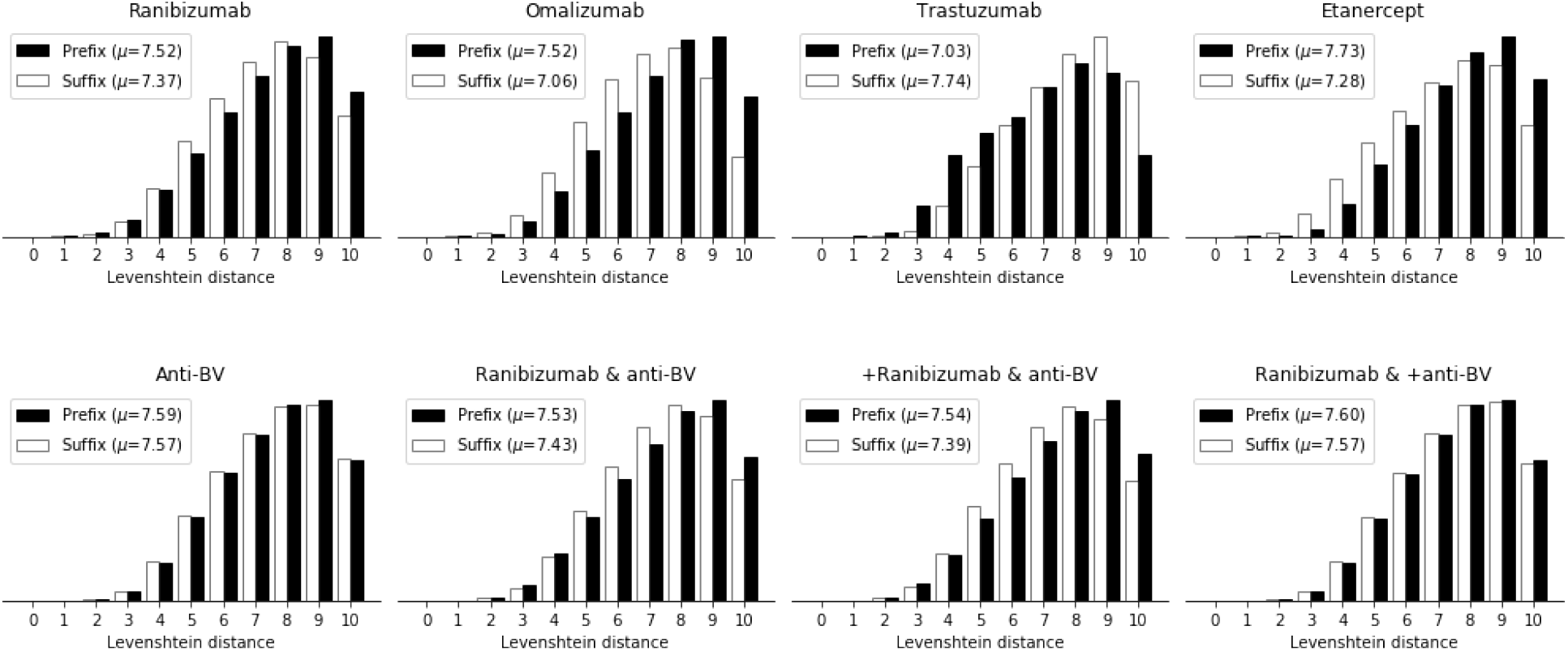
Prefix and suffix libraries have similar diversity. For each target evaluated, we compute the pairwise Levenshtein distances of the corresponding prefix (top) and suffix library (bottom).

#### E.3 Details on nonspecific motif enrichment analysis

For motif enrichment, we applied STREME motif enrichment analysis from the DREME suite to search for motifs enriched in FW-kappa over the BV library [21].For all STREME motif enrichment analysis, 490,000 sequences were sampled from both the primary and negative libraries and STREME was run with the following parameters: –protein –minw 3 –maxw 4 –nmotifs 100. After construction of the BV factorized library optimized for limited polyspecificity, we conducted motif enrichment analysis on each library to check whether known nonspecific motifs were decreased in the BV library when compared to the FW-kappa randomized library. As a preliminary analysis, we computed the number of known tryptophan, valine, arginine, and glycine nonspecific motifs (as identified in [9]) in a sample of 490,000 sequences in each library. (7). We report a few nonspecific motifs enriched in FW-kappa with significance values (Table 2). We observe that nonspecific motifs are enriched in the FW-kappa library over the BV factorizable library, further suggesting that the library has a favorable developability profile.

**Fig. 7.**
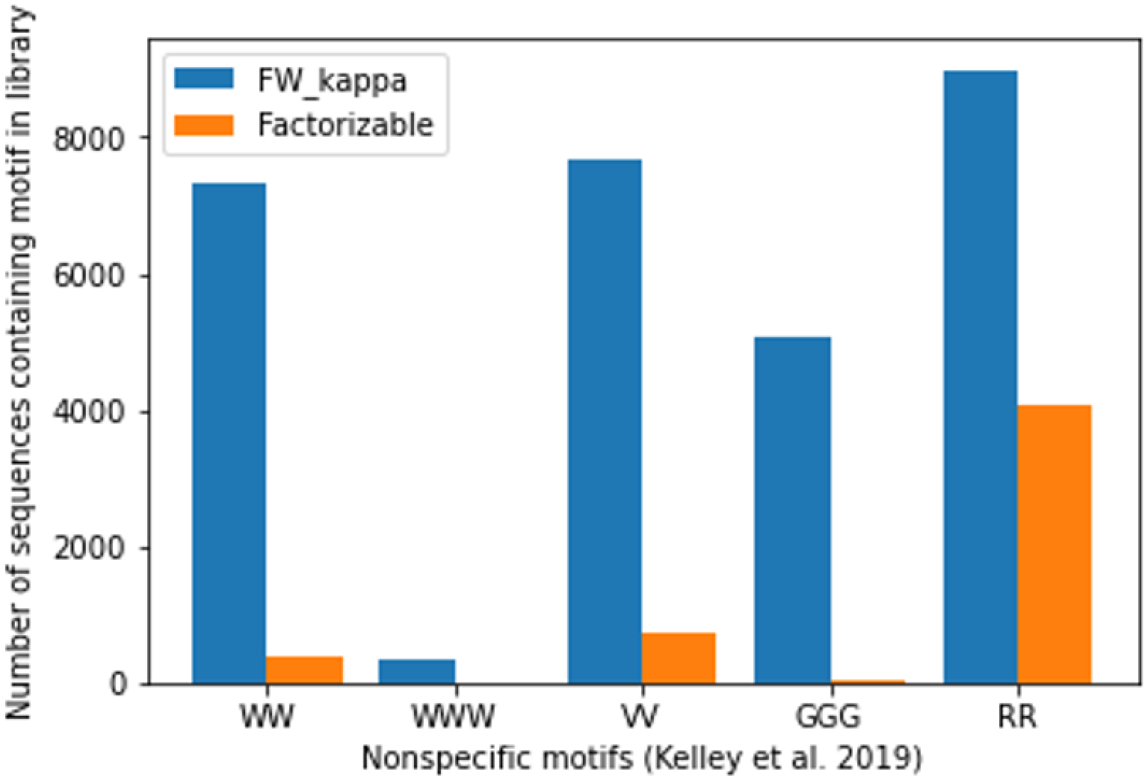
Ranibizumab Nonspecific motif counts in factorizable libraries vs. FW-kappa library

**Table 2.** Nonspecific motifs from STREME output for enriched sequences in FW-kappa over negative BV library

| Motif | p-value | no. of sites |
| --- | --- | --- |
| YYY | 3.3e-3456 | 159437 |
| GRG | 1.5e-1007 | 79625 |
| DVV | 6.3e-080 | 3959 |
| GGHS | 1.2e-054 | 3324 |
| GGD | 3.4e-028 | 1701 |
| WGG | 3.8e-014 | 434 |
| VGVD | 1.6e-013 | 622 |

## Notes

https://github.com/gifford-lab/FactorizableLibrary

